# Nomination of a novel plasma protein biomarker panel capable of classifying Alzheimer’s disease dementia with high accuracy in an African American cohort

**DOI:** 10.1101/2024.07.27.605373

**Authors:** Lindsey A. Kuchenbecker, Kevin J. Thompson, Cheyenne D. Hurst, Bianca M. Opdenbosch, Michael G. Heckman, Joseph S. Reddy, Thuy Nguyen, Heidi L. Casellas, Katie D. Sotelo, Delila J. Reddy, John A. Lucas, Gregory S. Day, Floyd B. Willis, Neill Graff-Radford, Nilufer Ertekin-Taner, Krishna R. Kalari, Minerva M. Carrasquillo

## Abstract

**Introduction:** African Americans (AA) are widely underrepresented in plasma biomarker studies for Alzheimer’s disease (AD) and current diagnostic biomarker candidates do not reflect the heterogeneity of AD.

**Methods:** Untargeted proteome measurements were obtained using the SomaScan 7k platform to identify novel plasma biomarkers for AD in a cohort of AA clinically diagnosed as AD dementia (n=183) or cognitively unimpaired (CU, n=145). Machine learning approaches were implemented to identify the set of plasma proteins that yields the best classification accuracy.

**Results:** A plasma protein panel achieved an area under the curve (AUC) of 0.91 to classify AD dementia vs CU. The reproducibility of this finding was observed in the ANMerge plasma and AMP-AD Diversity brain datasets (AUC=0.83; AUC=0.94).

**Discussion:** This study demonstrates the potential of biomarker discovery through untargeted plasma proteomics and machine learning approaches. Our findings also highlight the potential importance of the matrisome and cerebrovascular dysfunction in AD pathophysiology.

## Introduction

Until recently, a definitive diagnosis of Alzheimer’s disease (AD) was only achievable through post-mortem observation of key features of AD neuropathology in the brain, amyloid-β (Aβ) containing plaques and hyperphosphorylated tau tangles.^1^ Advancements in the ability to detect these markers ante-mortem through the use of cerebrospinal fluid (CSF) measurements of Aβ42/40, tau phosphorylated at threonine 181 (p-tau181), and measurement of amyloid pathology through positron electron tomography (PET) imaging has improved confidence in the diagnosis of AD.^2–4^ However, there is a need to develop diagnostic methods that are less costly and invasive than PET imaging and CSF measurement to improve access to accurate diagnostic information for the general population. Thus, the development of blood-based biomarkers is of major interest, owing to the potential of improving access to a confirmatory diagnosis for AD, and particularly for screening and monitoring of AD drug trials.^5^

Currently, several promising plasma biomarker candidates show high accuracy in identifying amyloid PET-positive individuals, such as Aβ42/40 (area under the curve [AUC]: 0.88),^6^ p-tau181 (AUC: 0.80-0.81),^7^ and p-tau217 (AUC: 0.92-0.96).^8^ While amyloid and tau are key components of AD pathophysiology, there is strong evidence of dysregulation in a wide array of biological pathways, such as neuroinflammation,^9^ synaptic function,^10^ lipid trafficking,^11^ and many others.^12,13^ Given the limited success of anti-amyloid therapies, it is apparent that these diverse processes may play an important role in AD risk and pathogenesis, contributing to the complexity and heterogeneity observed in individuals with AD.^14–16^ Thus, there is a need to identify novel biomarkers that represent disease processes beyond amyloid and tau.

The vast majority of AD biomarker studies have been performed in predominantly non-Hispanic White (NHW) cohorts. African American (AA) individuals have approximately double the risk of developing AD compared to NHW^17,18^ yet remain underrepresented in AD research.^19,20^ The few plasma biomarker studies that have been conducted in AA cohorts are limited in sample size, number of biomarkers tested, and often focus on differences in biomarker levels between NHW and AA.^21–25^ To date, only one published study has evaluated plasma p-tau217 in an AA cohort and reported that this marker achieved high accuracy in distinguishing autopsy-confirmed AD and control AA decedents (AUC=0.96, n=31), yet displayed lower accuracy in identifying AA study participants with clinical diagnoses of AD dementia (AUC=0.68, n=98).^21^

Given the well-established differences in AD prevalence^26,27^ and risk^28,29^ specific to this population, it is unclear if plasma proteins that have yet to be evaluated as biomarkers could improve diagnostic accuracy in this population. Genome-wide association studies have shown different frequencies and effect sizes of genetic variants in AD-related genes (*APOE, ABCA7, TREM2*) and biological pathways (lipid trafficking, immune regulation) in AA compared to NHW, indicating that there may be differential mechanisms driving AD pathogenesis in AA.^30–34^ This study aims to identify novel plasma protein biomarkers for AD specifically in AA. Here we combine untargeted proteomics and machine learning approaches to identify a novel set of plasma protein biomarkers capable of classifying AA study participants with clinical diagnoses of AD dementia and cognitively unimpaired (CU) with high accuracy. Additionally, this approach enables the investigation of differentially abundant proteins, pathways, and co-expression modules in plasma from AA study participants who have been diagnosed with AD dementia, providing insight into factors that drive or result from AD pathogenesis.

## Methods

### Cohort characteristics

This study includes 183 patients with AD dementia and 145 cognitively unimpaired elderly controls from the Florida African American Alzheimer’s Disease Studies (FCA^3^DS) cohort, who self-identified as AA and were enrolled in research studies of memory and aging at Mayo Clinic (Jacksonville, FL, USA; **Table 1**). Enrolled participants or their legally authorized representatives provided informed consent to participate in research specific to AD and related dementias, including the storage of biofluid samples for research use. Study participants underwent a standard clinical interview, neurological evaluation, and tests of memory and thinking as specified by individual study protocols. Alzheimer’s dementia diagnoses were established by a multidisciplinary panel referencing published criteria.^35,36^

**Table 1:** Cohort characteristics.

|  | **CU (n=145)** | **AD (n=183)** | **p-value** |
| --- | --- | --- | --- |
| Age at blood collection, y (mean ± SD) | 70.2 ± 7.93 | 77.3 ± 9.62 | <0.001 |
| Female sex, n (%) | 108 (74.5) | 129 (70.5) | 0.423 |
| *APOE*-ε4 carriers, n (%)<sup>*</sup> | 49 (33.8) | 106 (57.9) | <0.001 |

<sup>\*</sup>*APOE* genotype information available from 179/183 AD study participants. Statistical inference of continuous and categorical measures was performed using student t-test and chi-squared tests, respectively. Abbreviations: CU = cognitively unimpaired, AD = Alzheimer's dementia, *APOE* = apolipoprotein E gene.

### Plasma samples

Plasma samples from Mayo Clinic study participants included in this study were collected between January 1993 and July 2023. Blood samples were collected into EDTA Vacutainer tubes and promptly transported to the research laboratory. Blood was centrifuged at 2000 x g at 4°C for 10 minutes, aliquoted into 250 uL or 500 uL portions, and stored at -80°C until use. Plasma samples were thawed over ice for 30 minutes, gently agitated, aliquoted into 130 uL portions, randomized, and shipped to SomaLogic (Boulder, CO) for proteome measurements. Thus, plasma samples underwent only one freeze-thaw cycle prior to measurement.

### Genotyping

DNA was extracted from whole blood using the FLEX STAR system (AutoGen, MA, USA) and the Flex-iGene DNA Kit (Qiagen, Hilden, Germany) according to manufacturer’s protocols. TaqMan genotyping was performed to obtain *APOE* genotypes as described previously.^22^

### SomaScan untargeted proteomics

Plasma proteins were measured using the commercially available SomaScan 7k v4.1 assay (SomaLogic, Boulder, CO), a highly multiplexed, DNA aptamer-based assay that enables the measurement of more than 7,000 protein analytes with a 10^10^ dynamic range and minimal variability^37,38^ The SomaScan Assay v4.1 utilizes SOMAmer® (Slow Off-rate Modified Aptamer) reagents made up of short single-stranded DNA sequences modified with hydrophobic appendages to enable high-affinity binding to a diverse range of protein targets. Briefly, SOMAmer reagents form specific, high-affinity complexes with plasma proteins which are then quantified as relative fluorescent units (RFU) by hybridization to complementary sequences on DNA microarrays. Data standardization and quality control were performed by standard SomaLogic procedures, including hybridization normalization, intraplate median signal normalization, plate scaling, SOMAmer calibration, and adaptive normalization by maximum likelihood to adjust for variability across samples, SOMAmers, and batches.^37^ Protein levels were log2 transformed to approximate a normal distribution prior to downstream analyses.

### Validation of proteomics findings

Given that the SomaScan 1.1k assay includes 949 proteins that can also be detected with the 7k assay, we obtained a publicly available SomaScan 1.1k plasma proteome dataset from ANMerge^39,40^ European study participants with diagnoses of AD dementia or CU that were utilized as a replication dataset (**Supplementary Table 1**). In addition, the ability to replicate plasma protein findings in brain tissue was assessed using the Accelerating Medicine’s Partnership for AD (AMP-AD) Diversity Initiative dataset.^41^ Five AMP-AD data contributing sites provided brain tissue samples from their respective post-mortem cohorts, including Mayo Clinic, Rush University, Mount Sinai School of Medicine, Columbia University, and Emory University. This study utilized tandem mass tag coupled with mass spectrometry (TMT-MS) proteomics data generated by Emory University, containing 9,180 proteins quantified from dorsolateral prefrontal cortex (DLPFC) tissue from 517 AA, NHW, and Hispanic/Latino decedents that were autopsy-confirmed as AD or control (i.e. no neuropathology indicative of other neurodegenerative conditions; **Supplementary Table 2**).^42^ Receiver operating characteristic (ROC) analysis was performed using the pROC() R package to evaluate the classification accuracy of protein panels to distinguish between clinical (ANMerge, AD dementia vs. CU) and autopsy-confirmed diagnosis (AMP-AD, AD vs. control).

### Statistical analyses

#### Differential protein abundance (DPA) analysis

The association between individual plasma protein levels and diagnosis (AD dementia vs. CU) was evaluated using linear regression models, adjusting for technical and biological covariates: Model 1) log2(protein abundance) = Diagnosis + age at blood collection + sex + subarray, Model 2) log2(protein abundance) = Diagnosis + age at blood collection + sex + subarray + *APOE* genotype. The linear regression β-coefficients represent log2 fold change. Statistical significance was defined false discovery rate (FDR) adjusted p-values < 0.05 (q<0.05).^43^

#### Weighted gene co-expression network analysis (WGCNA)

Network analysis was performed using the Weighted Gene Correlation Network Analysis (WGCNA, v1.72-5; R version 4.3.2) algorithm on all 7298 plasma proteins for AD dementia and CU study participants.^44^ The WGCNA::bockwiseModules function was executed with a soft threshold power of 7.0, deepsplit of 4, minimum module size of 30, merge cut height of 0.07, mean topological overlap matrix (TOM) denominator, using biweight midcorrelation (bicor), signed network type, pamStage=TRUE, pamRespectsDendro=T, and a reassignment threshold of 0.05. This function creates a correlation matrix between protein pairs using bicor, which is then transformed into a signed adjacency matrix. The adjacency matrix is then used to calculate topological overlap (TOM), representing expression similarity across samples for all proteins in the network. Modules are identified using hierarchical clustering (as 1 minus TOM) and dynamic tree cutting. Following network construction, module eigenprotein (ME) values are defined by the first principal component of a given module. The MEs are considered representative protein abundance values for a module and explain modular protein covariance (citation). Module membership and determination of hub proteins within a given module was defined using Pearson correlation comparing proteins and MEs, defined as kME. Box plots represent the median, 25^th^ and 75^th^ percentile extremes, with the hinges of a box representing the interquartile range of the two middle quartiles of data within a group. Error bars were defined by the farthest data points up to 1.5 times the interquartile range away from the box hinges. Groupwise correlation significance was determined using student’s t-test and statistical significance was defined as p<0.05.

#### Gene Ontology (GO) analysis

Relevant gene ontology (GO) terms were determined for each protein module by gene set enrichment analysis (GSEA) using hypergeometric overlap with a Fisher’s exact test (FET). For enrichment, p<0.05 and a minimum of 5 genes per ontology must be present. A list of GO annotations from the Bader lab (formatted as a .GMT file and updated monthly)^45^ was used and implemented with an R script as previously published.^46^ Briefly, the script performs one-tailed FET using the R package Piano.

#### Brain cell type specificity

Cell-type enrichment was performed using a publicly available (GitHub - edammer/CellTypeFET: Calculate and Plot Significance of Gene List Overlap) cell-type marker list curated from Sharma et al.^47^ and Zhang et al.^48^ which contains 5 brain-specific cell-type marker lists (neurons, astrocytes, microglia, oligodendrocytes, and endothelia). The reference marker lists were generated by merging the Sharma and Zhang lists, therefore proteins that assigned to two cell types would default to that which was defined by Zhang et al to ensure each protein was only assigned to one cell type. Enrichment of cell types was determined by Fisher’s exact test and FDR correction (q<0.05).

#### Network preservation

The preservation of network modules between plasma and brain data was determined using the WGCNA::modulePreservation() function in R with the following parameters: 500 permutations, random seed set to 1 (for reproducibility) and quickCor set to 0. The plasma network was used as the test network and the brain as the reference network to calculate Zsummary preservation scores of plasma modules in brain.

### Predictive modeling

A median filter of greater than 7 was applied to log2 transformed SomaScan proteomics data to remove lowly abundant proteins. Multiple models were considered which included age at blood collection, sex, *APOE* genotype, and plasma protein predictors selected by machine learning. The data was centered and scaled prior to implementing an elastic net model, which was optimized using cross-validation with the nestedcv package (v 0.7.8)^49^ in R (Version 4.2.2). Five-fold cross-validation was performed in both the inner and outer loops. Five different feature selection approaches were evaluated, including t-test, signal-to-noise^2, variable importance, partial least squares, and the ReliefF algorithm. Feature sets of 10, 50, 100, and 250 were evaluated, and the AUC served as the performance metric. The nestedcv.train wrapper to the caret (version 6.0.94) package^50^ was then used to compare the nested cross-validation model performance of additional machine learning algorithms, including random forest, support vector machine, bayesian generalized linear model, mixture discriminant analysis, and several boosting methods (gradient boosting, extreme gradient boosting, bagged ADA boost, and logit boosting). Repeated nested cross-validation using 50 PLS-selected features was implemented on the same sample partitions to ensure a reliable estimate of each model’s performance. All other parameters were set to their default values.

## Results

### Cohort characteristics

Study participant characteristics are displayed in **Table 1**. Study participants diagnosed with AD dementia were slightly older than those diagnosed as CU. While the distribution of sex was consistent between CU and AD dementia, females are somewhat overrepresented, comprising 72.2% of the total cohort, compared to 61.8% of AA aged 65 and older in the population from which this cohort was drawn.^51^ As expected, the number of *APOE*-ε4 carriers was significantly larger in study participants diagnosed with AD dementia.

### Plasma proteins selected by machine learning predict AD dementia with high accuracy

Nested cross-validation was employed to optimize an elastic net approach to classify 328 AA participants using the 7298 human proteins that were quantified with the SomaScan 7k platform. Of the multiple feature selection methods and feature set sizes, the best AUC was observed for 50 features selected by partial least squares. Model stability was subsequently evaluated with 25 repeated nested cross-validation models, where the mean AUC was observed to be 0.90 (95% CI: 0.87-0.92; **Fig 1A**). During the optimization of 25 models, 50 plasma proteins were selected, including 33 proteins consistently observed across iterations. We evaluated the Shapley values of the 33 plasma protein predictors consistently identified, including two transgelin (TAGLN) measurements (**Fig 1B**). Both representations of TAGLN were among the least important features in the final model, yet they were both significantly more abundant in plasma samples from AD dementia cases compared to CU controls (**Fig 2**). The Shapley values, variable importance measures, and median abundance levels of the 33 plasma protein predictors are reported in **Supplementary Table 3**.

**Figure 1:**
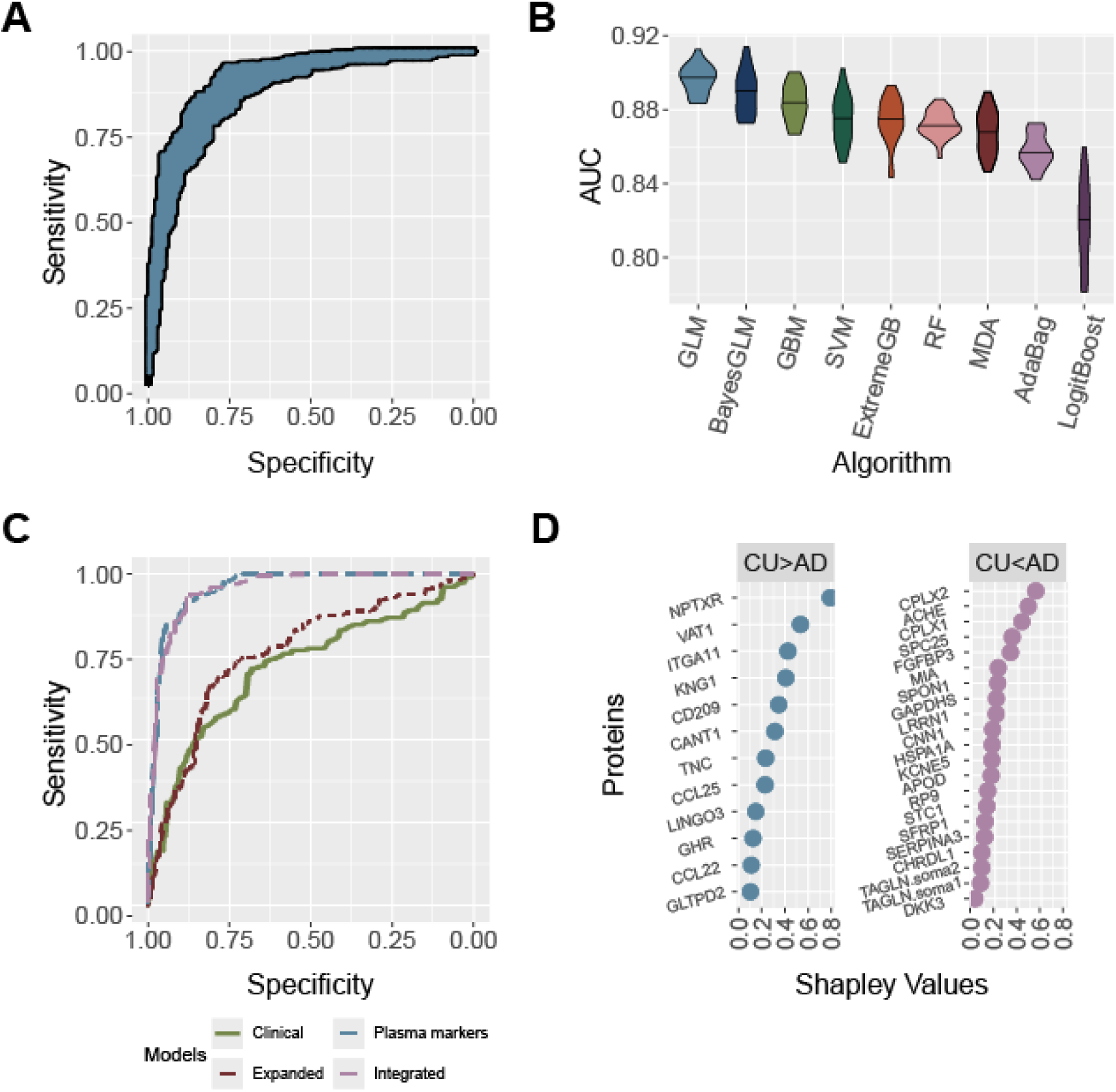
Classification efficacy of plasma proteins selected by machine learning approaches. **A)** Receiver operating characteristic (ROC) curve depicting the area under the curve (AUC) from repeated optimization of an elastic net model with partial least squares (PLS) selection of 50 features. **B)** Violin plots show the AUC of 9 machine algorithms tested to select the optimal model using the feature selection method shown in A. **C)** ROC curve shows the ability of a base clinical model (age at blood collection and sex; green), expanded clinical model (age at blood collection, sex, and *APOE* genotype; maroon), 33 plasma proteins selected through a machine learning model outlined in A and B (blue), and an integrated model including age, sex, *APOE*, and 33 plasma protein predictors (purple) to correctly classify AD dementia and CU study participants. **D**) Shapley values for the 33 plasma protein markers used in the predictive model. Proteins with decreased levels in AD dementia are shown in blue and proteins with increased levels in AD are shown in purple.

**Figure 2:**
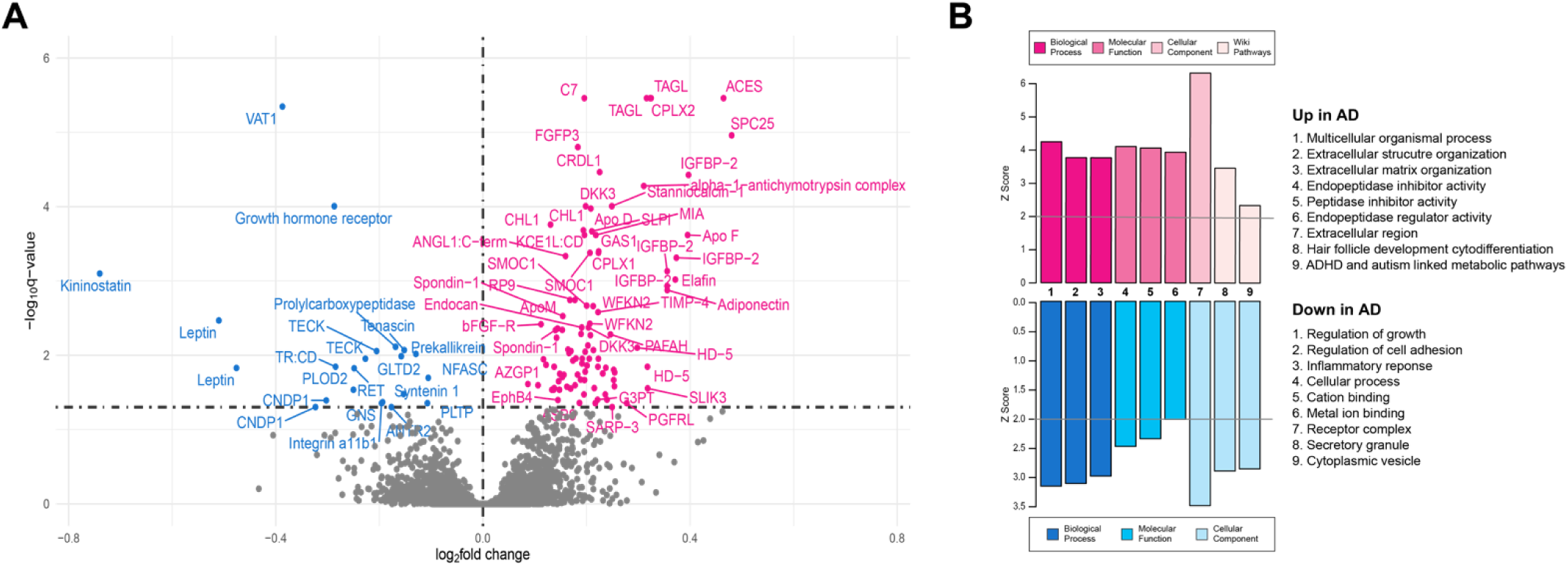
Differential abundance analysis comparing proteins associated with clinical diagnoses of AD dementia and CU. **A)** Volcano plot displaying the log2 fold change (x axis) and -log10 q-values (y axis) for plasma proteins differentially abundant in AD dementia compared to CU. Proteins with significantly increased levels (q<0.05) in AD dementia are shown in pink, proteins with significantly decreased levels in blue, and proteins that did not meet statistical significance (q>0.05) shown in grey. **B)** Top gene ontology (GO) terms associated with proteins significantly increased (pink) and decreased (blue) in AD dementia. GO libraries, from left to right, include biological process, molecular function, cellular component, and WikiPathways.

A base clinical model was developed using age at blood collection and sex, yielding an AUC of 0.67 (95% confidence interval [CI]: 0.65-0.67) An expanded clinical model including age at blood collection, sex, and *APOE* genotype showed improvement over the base clinical model (AUC=0.78 95%CI: 0.78-0.79). The optimized elastic net model of 33 protein markers yielded an AUC of 0.91 (95%CI: 0.91-0.91), representing a 24% increase in AUC from the base clinical model. Integrating the protein markers with age at blood collection, sex, and *APOE* genotype (integrated model) did not improve the model further (AUC=0.91, 95%CI: 0.90-0.91).

### Plasma proteomics reveals differentially altered markers in AD

Differential protein abundance analysis was performed to uncover altered levels of plasma proteins in AD dementia compared to CU. Differentially abundant proteins (DAPs) were defined by q-values < 0.05. As shown in **Fig 2A**, there were 98 proteins with significantly increased abundance and 22 proteins with significantly decreased abundance in AD dementia. A sensitivity analysis additionally adjusting for *APOE* genotype uncovered 84 DAPs, 78 of which overlapped with the *APOE*-unadjusted model. Differentially abundant proteins include several markers previously identified in AD including, but not limited to, insulin-like growth factor-binding protein 2 (IGFBP-2; q=3.47×10^-6^) SPARC-related modular calcium binding 1 (SMOC1; q=4.19×10^-4^), and neural cell adhesion model 1 (NCAM1; q=4.59×10^-3^). Pathway analysis of the 98 proteins with significantly higher levels in AD dementia revealed enriched GO terms related to the matrisome and peptidase signaling. Enriched pathways in the 22 proteins with significantly lower levels in AD dementia were associated with regulation of cell adhesion, growth, and receptor complexes.

### Plasma protein co-expression modules associate with AD demenita

Weighted correlation network analysis was performed to characterize protein communities among the 7298 plasma proteins using the WGCNA algorithm. There were 12 co-expression modules identified that ranged in size from 1668 protein members (M1) to 96 members (M12) (**Fig 3A**). The relationship between plasma modules and key covariates was examined using weighted correlation between MEs and age at blood collection, sex, *APOE* genotype, and AD dementia clinical diagnosis. There were 14 significant correlations (p<0.05) between modules and covariates. GSEA was performed to identify representative biological domains within modules (**Supplementary Table 4**) and top representative terms were selected to annotate the modules. Enrichment for cell type markers from 5 brain cell types was also performed to assess whether proteins previously isolated from brain cell types could be captured in our plasma modules. There were 8 significant (p<0.05) module associations with cell type markers across the 12 plasma modules and each of the 5 cell types had at least 1 significant result (**Fig 3A**).

**Figure 3:**
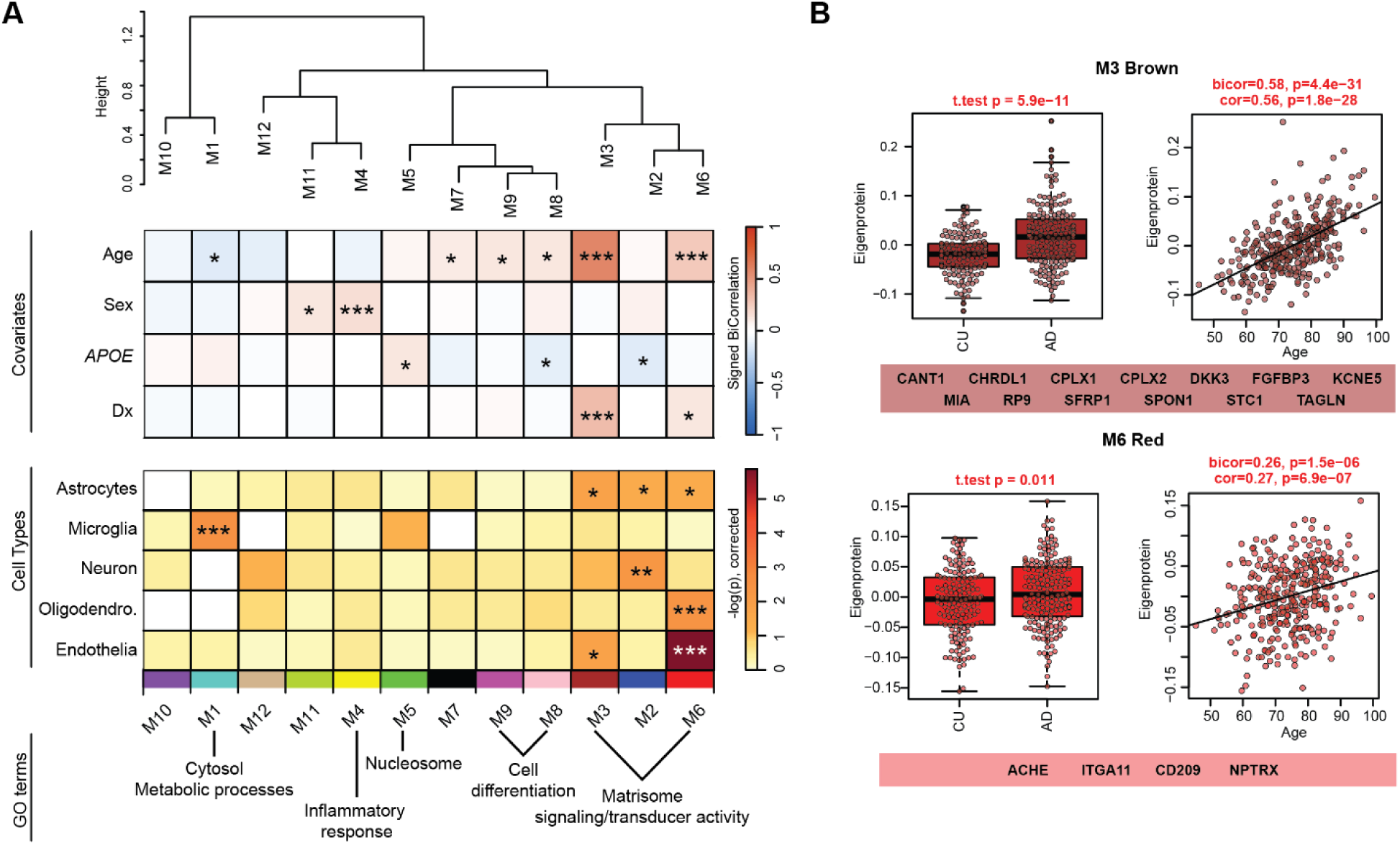
**A)** Weighted gene correlation network analysis (WGCNA) was performed with 7298 proteins measured from plasma and yielded 12 co-expression modules. A hierarchical clustering dendrogram illustrates module relatedness within the network based on module eigenprotein (MEs) values. The correlation (bicor) of MEs with sample covariates age, sex, *APOE* genotype, and diagnosis (Dx) is visualized as a heatmap (red indicated positive correlation and blue indicated negative correlation). The unweighted heatmap depicts enrichment of brain cell type markers (determined by one-way Fisher’s exact test) across the plasma modules for astrocyte, microglia, neuron, oligodendrocyte and endothelial cell types (darker color indicates stronger enrichment). Representative GO terms for select modules, based on gene set enrichment analysis are also shown. **B)** MEs grouped by diagnosis (CU and AD) were plotted as box and whisker plots for modules M3 (brown) and M6 (red), chosen based on significant association with AD dementia diagnosis. Group-wise MEs were compared using student t-test, unadjusted p-values are shown (M3: p=5.9E-11; M6: p=0.01). Box plots represent median, 25^th^ and 75^th^ percentiles. Box hinges represent the interquartile range from the box hinge. Correlation of MEs and age are shown as scatterplots, bicor and Pearson correlation with associated p-values are shown (M3: bicor=0.58, p=4E-31; cor=0.56, p=1.8E-28. M6: bicor=0.26, p=1.5E-06; cor=0.27, p=6.9E-07). Plasma protein predictors selected by machine learning that are members of modules M3 and M6 are shown under the box and scatter plots. Heatmap statistical significance is defined as 0.05>p>0.01 = *, 0.01>p>0.005 = **, p<0.005 = ***.

**Figure 4:**
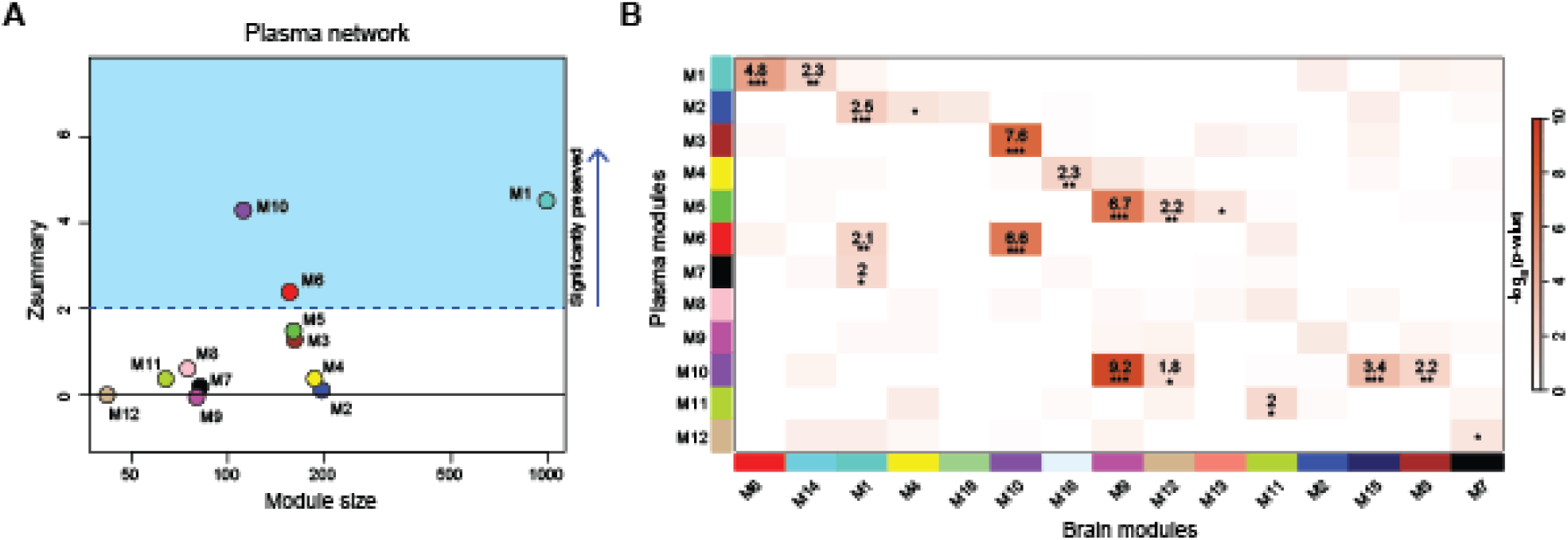
Assessment of module preservation between plasma and brain protein networks. **A)** module preservation was performed for the 12 plasma protein co-expression modules in 18 brain co- expression modules. Significant preservation was determined by modules with a Zsummary ≥1.96 (p = 0.05, dashed blue line). **B)** Overrepresentation analysis (ORA) was employed to assess the overlap of plasma module members with brain module members and the heatmap was generated to show enrichment across networks. Significant module overlap was determined by one-tailed Fisher’s exact test and -log10 p-values are displayed with accompanying significance indicated as stars (p<0.05 = *, p<0.01 = **, p<0.001 = ***), where darker red color indicates stronger enrichment.

There were 2 modules significantly correlated with AD dementia, M3 (brown) and M6 (red) (**Fig 3A**). Eigenprotein values for M3 and M6 were significantly higher in AD dementia compared to CU and significantly positively correlated with age (**Fig 3B**). Astrocyte and endothelia cell type markers were significantly enriched in both M3 and M6, with the addition of significant enrichment for oligodendrocyte markers in M6 (**Fig 3A**). Both modules M3 and M6 shared multiple top enriched GO terms primarily related to the matrisome, including cell adhesion and extracellular matrix (**Fig 3A, Supplementary Table 4**). Module membership of the 33 predictive plasma proteins was then considered. There were 14 predictive proteins in M3 and 4 predictive proteins in M6.

### Network preservation analysis of plasma and brain co-expression modules

Module preservation analysis using the plasma network as the test dataset and the brain as the reference was performed to determine which of the 12 plasma modules (**Supplementary Table 5**) were preserved with the 18 modules from the brain network (**Supplementary Table 6**) generated from the AMP-AD dataset. Of the 12 plasma modules, 3 were significantly preserved in brain (Zsummary>1.96 or p≤0.05). Overrepresentation analysis (ORA) was also performed to determine the enrichment of overlapping proteins from the plasma modules in the brain modules. The ORA results demonstrate greater overlap across networks as compared to the network preservation analysis, with 10 out of the 12 plasma modules significantly represented in the brain modules. AD-related plasma modules of interest (M3 and M6) were among the top significantly represented in brain modules. Only two plasma modules remained unrepresented in brain modules (M8 and M9) which were enriched for GO terms related to cell differentiation. Notably, several plasma modules were significantly enriched across multiple brain modules, which may explain the lack of module preservation for most plasma modules in brain as preservation is based on measures of topology and connectivity within modules, whereas ORA is based on enrichment of protein module members. Collectively, these results indicate that plasma proteins are well represented in brain-derived modules, however, the network structure is markedly different in each tissue type. As expected, plasma protein modules also contain non-overlapping features with brain network modules.

### Validation of plasma protein predictors in external, non-overlapping datasets

The reproducibility of the classification accuracy of plasma protein predictors identified through machine learning in the FCA^3^DS dataset was assessed in external, non-overlapping, plasma and brain datasets. Of the 32 unique plasma protein predictors selected through machine learning approaches in the FCA^3^DS cohort, 16 were present in the ANMerge SomaScan 1.1k dataset and 21 were present in the AMP-AD TMT-MS dataset **(Fig 5A).** Seven proteins were missing >20% data in the AMP-AD dataset and were thus excluded from downstream analysis.

**Figure 5:**
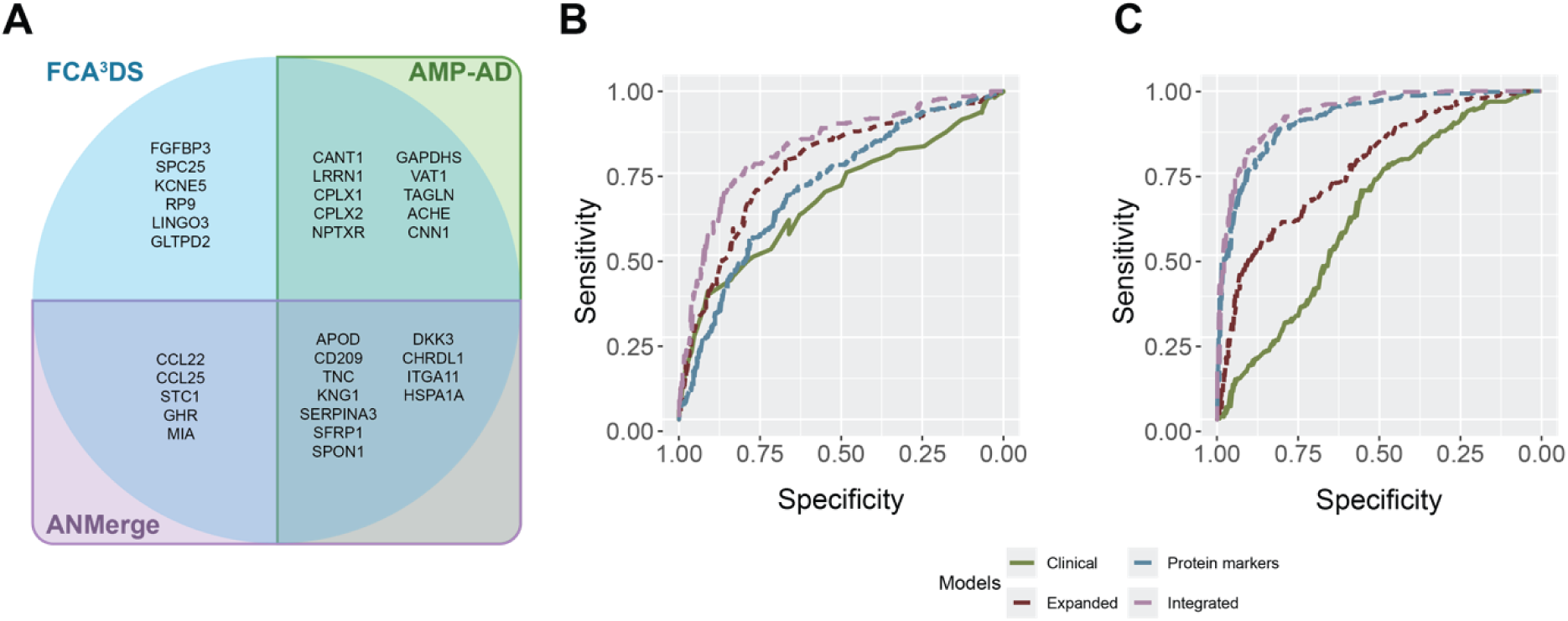
Replication of plasma protein predictors in external datasets. **A)** Venn diagram depicts 32 unique plasma protein predictors identified in the FCA^3^DS cohort (blue circle). The overlap of proteins present in external datasets are enclosed by rectangles, with ANMerge in purple and AMP-AD in green. Of these 32 proteins, 6 were present only in the FCA^3^DS cohort (upper left quadrant), 5 in FCA^3^DS and ANMerge (lower left quadrant), 10 in FCA^3^DS and AMP-AD (upper right quadrant), and 11 in all three cohorts (lower right quadrant). **B)** Receiver operating characteristic (ROC) analysis displays the accuracy of four models to distinguish between study participants with diagnoses of AD dementia and CU, including a base clinical model (age at blood collection and sex; green), expanded clinical model (age at blood collection, sex, and *APOE* genotype; maroon), plasma protein predictors (blue), and an integrated model including age, sex, *APOE*, and 16 plasma protein predictors that were present in the ANMerge SomaScan 1.1k dataset (purple). **C)** ROC analysis shows the accuracy of four models, as described for panel B, to distinguish between autopsy-confirmed diagnoses of AD and control. Of the 32 plasma protein predictors identified in the FCA^3^DS cohort, 14 plasma protein predictors with <20% missing data were present in the AMP-AD dataset and included in the ROC analysis.

For the ANMerge dataset, a base clinical model to classify 180 cognitively unimpaired study participants and 303 study participants with a clinical diagnosis of AD dementia was implemented using age at blood collection and sex, yielding an AUC of 0.67 (95%CI: 0.62-0.72). An expanded clinical model including age, sex, and *APOE* genotype improved classification performance by 10% (AUC=0.77, 95%CI: 0.73-0.82). A model including 16 plasma protein predictors (SFRP1, HSPA1A, KNG1, ITGA11, CD209, CCL22, MIA, CCL25, TNC, GHR, CHRDL1, DKK3, SERPINA3, SPON1, STC1, APOD) resulted in an AUC of 0.71 (95%CI: 0.66-0.76). Integrating the plasma markers with age, sex, and *APOE* genotype resulted in an AUC of 0.83 (95%CI: 0.79-0.87; **Fig 5B**).

In the AMP-AD dataset, a base clinical model to classify autopsy-confirmed controls (n=144) and AD brains (n=377), including age at death and sex, yielded an AUC of 0.51 (95%CI: 0.46-0.56). The addition of *APOE* genotype to the model increased the AUC to 0.72 (95%CI: 0.67-0.76). A model including 14 proteins isolated from DLPFC tissue (TAGLN, HSPA1A, CPLX2, CNN1, VAT1, CPLX1, KNG1, ITGA11, DKK3, SERPINA3, SPON1, GAPDHS, APOD, NPTXR) resulted in an AUC of 0.92 (95%CI: 0.89-0.94), a 20% increase over the model with age, sex, and *APOE*. Combining the protein predictors with age, sex, and *APOE* genotype yielded an AUC of 0.94 (95%CI: 0.91-0.96; **Fig 5C**). A sensitivity analysis including only AA decedents testing the plasma proteins integrated with age, sex, and *APOE* genotype achieved an AUC of 0.98 (95%CI: 0.96-0.99). In this sensitivity analysis, the 14 plasma proteins alone yielded an AUC of 0.94 (95%CI: 0.90-0.99; **Supplementary Table 7**)

## Discussion

In this study, a data-driven approach to biomarker discovery through untargeted proteomics and machine learning identified a novel plasma biomarker panel capable of predicting AD dementia with high diagnostic accuracy in an African American cohort (AUC=0.91). A subset of 14 proteins in this predictive panel are present in the AMP-AD Diverse Cohorts TMT-MS brain proteomics dataset and achieved an AUC of 0.92 to distinguish between decedents autopsy-confirmed as AD or control. Differential protein abundance analysis in plasma revealed 98 proteins with higher levels and 22 proteins with lower levels in ante-mortem study participants with AD dementia compared to CU controls. Network analysis uncovered 12 protein co-expression modules, two of which were significantly associated with clinical diagnosis and age. The disease-associated modules were found to be enriched with astrocytes and endothelial cells and enriched for GO terms related to the matrisome.

### Relevance of plasma protein predictors to AD pathophysiology

In support of the reproducibility and relevance of the findings presented in this study, many of the plasma protein predictors selected by machine learning have been previously reported to be involved in various aspects of AD pathophysiology. Of the 32 unique plasma protein predictors, 12 have been linked to hallmark AD pathology, including amyloid and tau. For example, SFRP1 and SPON1 are secreted proteins that impact amyloid precursor protein processing and can modulate production of pathogenic forms of Aβ and plaque formation.^52–54^ In addition to core AD pathology, 14 of the 32 proteins align with other biological processes previously shown to be disrupted in AD, including synaptic integrity and neuroinflammation. In particular, NPTXR is a presynaptic protein that shows persistent loss in proportion with the progression of AD in CSF.^55^ Similarly, the direction of change observed in plasma protein predictors identified in this study (including those mentioned above: SFRP1, SPON1 and NPTXR) demonstrate concordance with previous reports in brain, plasma, and CSF for those with existing measurements.^46,52,54,56–58^

At the level of biological pathways, plasma protein predictors identified in this study also align well with AD diagnosis and disease mechanisms according to network analysis. The greatest representation of plasma protein predictors was within the age and AD dementia-associated modules, M3 (brown) and M6 (red). Importantly, these AD dementia-associated modules were enriched for GO terms related to matrisome biology, including extracellular matrix and cell adhesion, which are consistent with previous proteomic results from human brain tissue.^59^ In brain, the “matrisome” module was significantly correlated with AD pathology and cognitive function and the protein members of this module have been hypothesized to represent potential biomarkers and therapeutic targets for AD as possible modulators of pathology and downstream consequences.^59^ While many of the predictive proteins nominated here have been previously linked to AD mechanisms, some proteins are novel markers not comprehensively studied in AD. This could be representative of the general heterogeneity of AD, the innovative combination of techniques applied in this study, and/or signatures specifically vulnerable in an understudied, diverse cohort.

### AD pathophysiology reflected in the periphery

Broad changes in structural and functional integrity of cerebrovasculature are commonly observed in AD and have been linked to neuroinflammatory responses, blood brain barrier (BBB) breakdown and worsening dementia.^60–63^ The extent to which centrally linked mechanisms, such as vascular dysfunction, can be captured in peripheral biofluids is not well-understood. In the present study, enrichment analyses were conducted to inform the biological relevance of potentially informative co-expression modules and address the unanswered question of the ability of plasma proteins to reflect perturbed pathways in AD. Importantly, AD dementia-related modules M3 and M6 are both enriched for endothelia and astrocytes, critical cellular constituents of vascular compartments. These modules are also enriched in pathways related to the matrisome, which have been linked to cerebrovascular dysfunction in AD.^64,65^ Together these findings may reflect AD-related changes in the vasculature and highlight the potential to capture central AD pathophysiology in a peripheral tissue. This is particularly relevant given the greater prevalence of vascular dysfunction in the AA population, highlighting the importance of biomarker development that accounts for heterogeneity across populations.^66,67^

### Study limitations and strengths

This study is not without limitations. There was limited availability of gold-standard diagnostic biomarkers such as amyloid-PET or CSF Aβ42/40. As a result, it was not possible to build models to assess the ability of plasma predictors to predict abnormal amyloid levels as measured through PET imaging or CSF. This study relied on clinical diagnosis as an outcome measure rather than autopsy-confirmed diagnosis. However, the strong predictive ability of a subset of identified plasma protein predictors in a post-mortem racially diverse replication cohort indicates that these proteins can distinguish between autopsy-confirmed AD and controls. Moreover, core AD plasma biomarkers such as p-tau217 and Aβ42/40 are not included in the SomaScan 7k assay and we were thus unable to assess the combined ability of these markers with plasma protein predictors to classify AD dementia and CU study participants. Future work is planned to measure core AD biomarkers in this cohort.

In summary, this study nominates a novel plasma protein biomarker panel capable of classifying AD dementia with high accuracy in an African American cohort. This work also highlighted molecular signatures and processes that potentially implicate cerebrovascular dysfunction as a key component of AD in this African American cohort. Assessing the generalizability of the predictive biomarker panel as well as individual and protein module associations in larger NHW and racially diverse cohorts is a critical next step to better understand the similarities and differences in AD pathophysiology across populations.

## Author contributions

Conceptualization: MMC, LAK

Funding acquisition: MMC

Investigation: MMC, LAK, KJT, CDH, BMO, KRK.

Methodology: MMC, LAK, KRK, KJT, CDH, MGH, JSR, TN, HLC, KDS, DJR

Project administration: MMC

Resources: MMC, JL, GSD, FBW, NGR, NET

Visualization: LAK, KJT, CDH

Writing – original draft: MMC, LAK, KRK, KJT, CDH, BMO

Writing – review and editing: all authors read and approved the final version of the manuscript.

## Acknowledgements

We thank the participants and their families for their participation in research. Without them this work would not have been possible. This work was funded by the Carl Angus DeSantis Foundation and supported by NIH/NIA U19 AG074879-01 and the Mayo Clinic Alzheimer’s Disease Research Center (P30-AG062677). The results published here are in part based on data obtained from the AD Knowledge Portal (https://adknowledgeportal.org/). Data generation was supported by the following NIH grants: U01AG046139, U01AG046170, U01AG061357, U01AG061356, U01AG061359, and R01AG067025. We thank the participants of participants of the Religious Order Study, Memory and Aging Project, the Minority Aging Research Study, Rush Alzheimer’s Disease Research Center, Mount Sinai/JJ Peters VA Medical Center NIH Brain and Tissue Repository, National Institute of Mental Health Human Brain Collection Core (NIMH HBCC), Mayo Clinic Brain Bank, Sun Health Research Institute Brain and Body Donation Program, Goizueta Alzheimer’s Disease Research Center, New York Brain Bank at Columbia University, New York Genome Center and the Biggs Institute Brain Bank for their generous donations. Data and analysis contributing investigators include Nilüfer Ertekin-Taner, Minerva Carrasquillo, Mariet Allen (Mayo Clinic, Jacksonville, FL), David Bennett, Lisa Barnes (Rush University), Philip De Jager, Vilas Menon (Columbia University), Bin Zhang, Vahram Haroutanian (Icahn School of Medicine at Mount Sinai), Allan Levey, Nick Seyfried (Emory University), Rima Kaddurah-Daouk (Duke University), Steve Finkbeiner (University of California-San Francisco/Gladstone Institutes), Daifeng Wang (University of Wisconsin-Madison), Stefano Marenco (NIMH HBCC), Anna Greenwood, Abby Vander Linden, Laura Heath, William Poehlman (Sage Bionetworks).

## Consent Statement

Written informed consent was obtained from all participants.

